# A dual motif mediates outer-membrane translocation and packing of glycosidases into *Bacteroides* Outer Membrane Vesicles

**DOI:** 10.1101/377861

**Authors:** Ezequiel Valguarnera, Nichollas Scott, Mario F. Feldman

## Abstract

Outer membrane vesicles (OMV) are spherical structures derived from the outer membrane (OM) of Gram-negative bacteria. *Bacteroides spp. are* prominent components of the human gut microbiota, and OMV produced with these species are proposed to play key roles in gut homeostasis. OMV biogenesis in *Bacteroides* is a poorly understood process. Here, we revisited the protein composition of *B. theta* OMVs by mass spectrometry. We confirmed that OMVs produced by this organism contain large quantities of glycosidases and proteases, with most of them being lipoproteins. We found that most of these OMV-enriched lipoproteins are encoded by polysaccharide utilization loci (PULs), such as the *sus* operon. We examined the subcellular localization of the components of the Sus system, and found that the alpha-amylase SusG is highly enriched in OMVs while the oligosaccharide importer SusC remains mostly in the OM. We show that all OMV-enriched lipoproteins possess a lipoprotein export sequence (LES) that mediates translocation of SusG from the periplasmic face of the OM towards the extracellular milieu and is required for SusG to localize preferentially to OMVs. We also show that surface-exposed SusG in OMVs is active and can rescue growth of bacterial cells incapable of growing on starch as only carbon source. Our results support the role of OMVs as “public goods” that can be utilized by other organisms with different metabolic capabilities.

**IMPORTANCE:** Species from the *Bacteroides* genus are predominant members of the human gut microbiota. OMVs in *Bacteroides* have been shown to be important for the homeostasis of complex host-commensal relationships, mainly involving immune tolerance and protection from disease. OMVs carry many enzymatic activities involved in the cleavage of complex polysaccharides and have been proposed as public goods that can provide growth to other bacterial species by release of polysaccharide breakdown products into the gut lumen. Nevertheless, the mechanistic nature of OMV biogenesis is unclear for *Bacteroides* spp. This works shows the presence of a negatively-charged rich amino acid dual motif that is required for efficient packing of the surface-exposed alpha-amylase SusG into OMVs. Discovery of this motif (LES) is the first step in the generation of tailor made probiotic interventions that can exploit LES-related sequences to generate *Bacteroides* strains displaying proteins of interest in OMVs.

## INTRODUCTION

Outer membrane vesicles (OMVs) are small spherical structures derived from the outer membrane (OM) of all Gram-negative bacterial species. OMVs are composed of phospholipids, lipopolysaccharide (LPS) or lipooligosaccharide (LOS), and OM and periplasmic proteins (1, 2). OMVs can mediate host-microbe interactions by facilitating long-distance delivery of virulence factors, modulating the host-immune response, and contributing to antibiotic resistance (1, 3–8). Despite these key roles in bacterial physiology, OMV biology is poorly understood.Recent research indicates that OMVs are produced by diverse mechanisms. For example, OMVs are proposed to be generated by LPS remodeling in *Porphyromonas gingivalis*, *Salmonella enterica* and *Pseudomonas aeuroginosa* (9-12). In contrast, OMVs from *Haemophilus influenza* and *Vibrio cholerae* are thought to be the result of an accumulation of phospholipids in the OM outer leaflet mediated by their specialized VacJ/Yrb transporter (13). Thus, it appears that there is not a universal mechanism of OMV biogenesis. For most species, including *Bacteroides* spp., OMV biogenesis remains poorly understood (5).

Species from the phylum Bacteroidetes compose a major part of the human gut microbiota, and OMVs from these organisms are proposed to play important roles in the commensal-host relationship (14, 15). These roles include the delivery of immunomodulatory molecules to host immune cells, an interaction that appears to help prevent colitis flares in patients with inflammatory bowel disease (IBD) (7, 8). Furthermore, *Bacteroides* OMVs have been proposed to interfere with intracellular Ca^2+^ signaling in host cells (16). Most studies focus primarily on two predominant species in the human gut, *B. thetaiotaomicron (“B. theta”) and B. fragilis*. These species produce large amounts of uniformly sized OMVs that have a distinct protein composition compared to the OM, indicating that these OMV particles are not by-products of bacterial lysis (5). Most *Bacteroides* OMV-exclusive proteins are putative acidic lipoprotein hydrolases, suggesting that proteins with similar structural and physicochemical properties are selectively sorted to OMVs (5). Many *Bacteroides* enzymatic lipoproteins are encoded on polysaccharide utilization loci (PULs), which constitute ~20% of the *B. theta* genome and are essential for the breakdown and acquisition of plant, fungi and mucosal complex polysaccharides (17–19). PULs consist of at least one TonB-dependent receptor similar to a SusC-like protein; one nutrient binding accessory protein or SusD; and other accessory binding proteins (20). PULs can also carry two-component systems that sense nutrient variations in the media, with subsequent induction of PUL genes required for the utilization of complex carbon sources (21). Hence, *Bacteroides* cells can modify the enzymatic composition of OMVs according to available carbon sources (5, 22). The enzymatic arsenal carried by *Bacteroides* OMVs appears to carry a “social” function, as the products of OMV-mediated hydrolysis can be utilized by other bacteria within the gut (22, 23).

Here, we further characterized the protein composition of *B. theta* OMVs. We confirmed that OMVs produced by this organism mainly contain putative acidic lipoprotein hydrolases or carbohydrate-binding proteins. Many of these OMV-enriched lipoproteins were found to be encoded by PULs. We examined the subcellular localization of the components of the archetypical PUL, the starch utilization system (Sus) (20, 24), and found that the alpha-amylase SusG and other Sus lipoproteins are highly enriched in OMVs. In contrast, the oligosaccharide importer SusC remains mostly in the OM. We show that the presence of a lipoprotein export sequence (LES) mediates translocation of SusG from the periplasmic face onto the extracellular milieu and is both required and sufficient for SusG to localize preferentially to OMVs. Our results support the role of OMVs as “public goods” that can be utilized by other organisms with different metabolic capabilities.

## RESULTS

### Proteomics analysis of membrane and OMV fractions from *B. theta*

Electron micrograph analysis confirmed that *B. theta* produces large amounts of uniform OMV particles (Figure 1). We previously performed a proteomic analysis of *B. theta* OM and OMV employing Triton X-100 for the purification of OM proteins (5). In this work, we employed N-lauroylsarcosine (Sarkosyl), which has been widely used for the separation of inner membrane (IM) and OM fractions in Gram-negative bacteria (25-27) to verify that the apparent selective fractionation of proteins into OM and OMV was not due to the use of a specific detergent. Using early stationary cultures from *B. theta,* we prepared the different membrane fractions as indicated in Figure S1. In our previous study, OMVs were not treated with the detergent employed to extract the IM proteins from total membrane preparations. To rule out possible detergent effects, we added an additional step consisting of incubating OMV in 1% Sarkosyl prior to the ultracentrifugation of the samples to recover an OMV supernatant (OMV-S) and an OMV pellet (OMV-P). Samples were lyophilized for proteomic analysis by LC-MS/MS and aliquots were visualized by Coomassie Blue (Figure S1).

**Figure 1.**
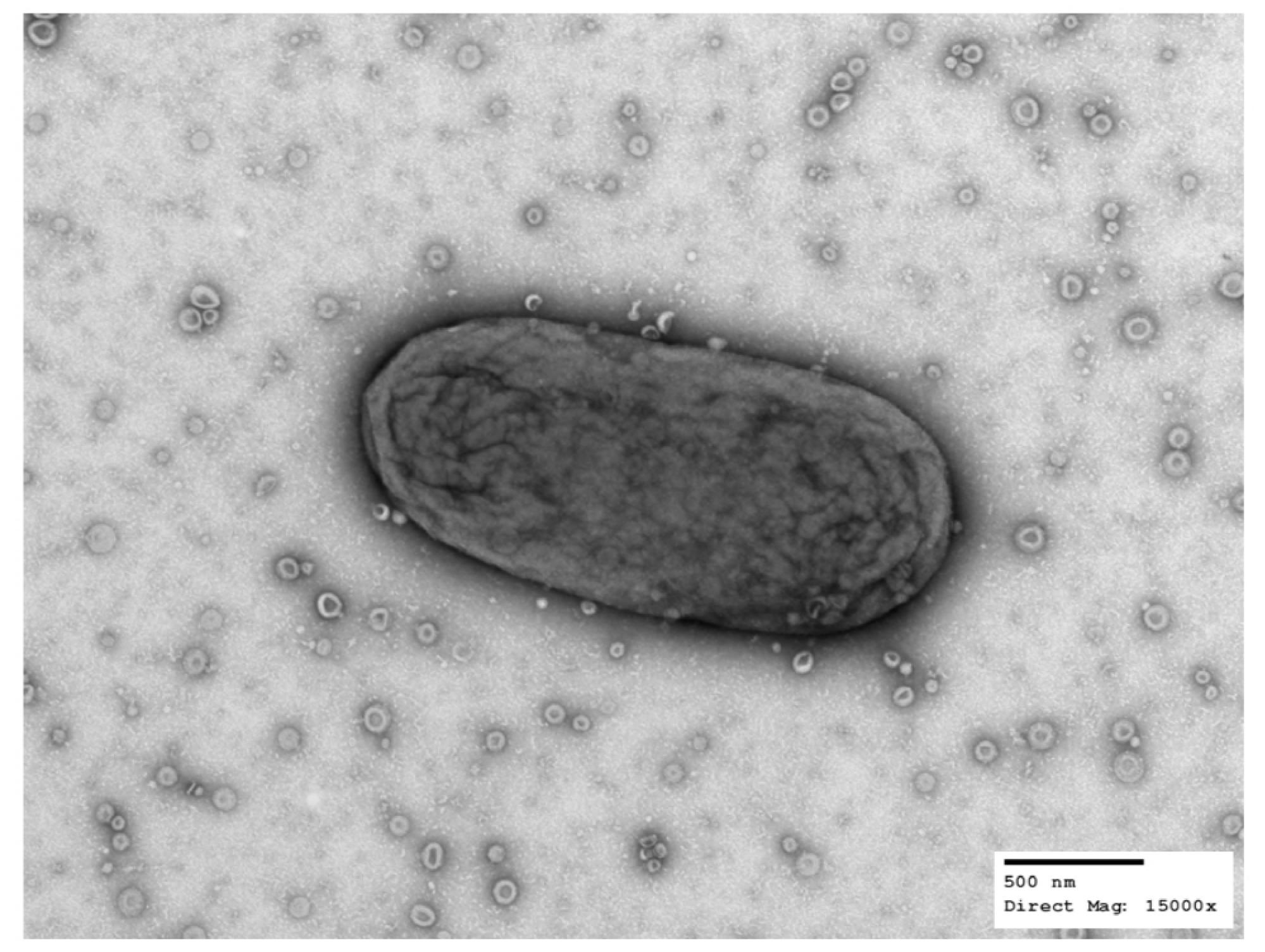
*B. theta* produces outer membrane vesicles (OMV). Transmission electron microscopy of a single *B. theta* cell together with OMVs in the extracellular milieu. *B. theta* cells were swabbed from a solid media plate, suspended in PBS and processed for TEM. Images were acquired at a direct magnification of 15000x. Scale bar represents 500 nm. Credit to Wandy Beatty, Molecular Microbiology Imaging Facility, WUSTL.

Our new dataset with the annotation and predicted function and localization for proteins in all fractions is provided as Table S1. We confirmed our previous findings showing an enrichment of lipoproteins in the OMV in comparison with the OM fraction (Table 1, Table S2) (5). Figure 2 highlights the top enriched proteins in the OMV. We found that 18 out of the 23 OMV-exclusive lipoproteins from our previous study are also enriched in the new OMV preparations (5). We then cloned three OMV and three OM-enriched proteins identified in our studies into the expression vector pFD340 as 6xHis-tagged proteins, to confirm their localization by Western Blot. Figure 3a shows that a putative cell surface protein (BT_1488), as well as a putative calpain-like protease (BT_3960), and a putative zinc peptidase (BT_3237), are highly enriched in OMVs. Figure 3b shows that BT_0418, BT_2844 and BT_2817, identified as OM-enriched proteins by MS, are retained at the OM and are not present in OMVs. BT_0418 is a Porin F ortholog (8 β-strand protein with a PG-binding domain), BT_2817 is a TonB-dependent receptor (22 β-strand protein), and BT_2844, a lipoprotein containing tetratricopeptide repeat (TPR) motifs. These set of experiments demonstrate that lipoproteins are indeed differentially sorted between OM and OMV.

**Figure 2.**
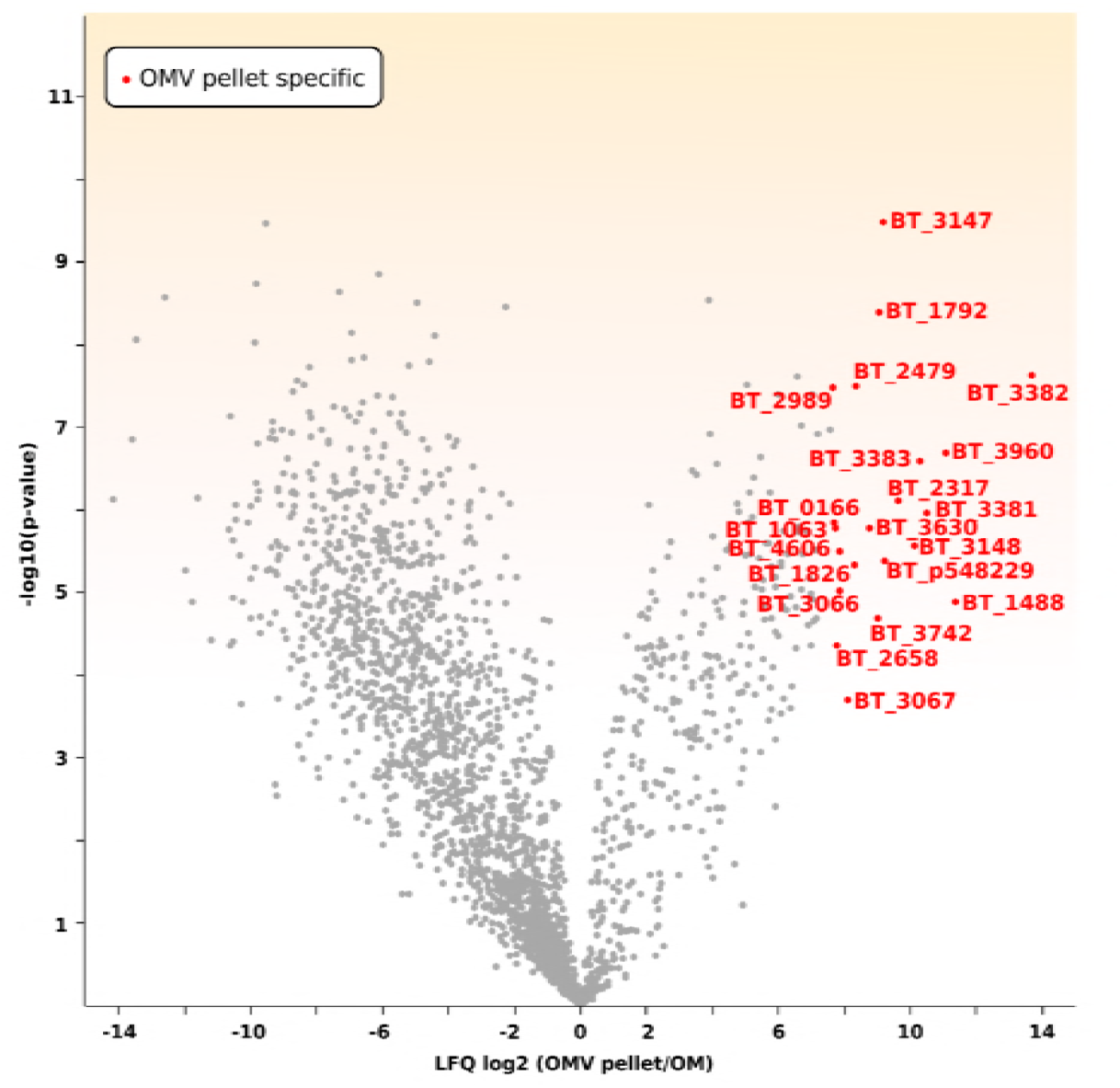
*B. theta* OMV protein content is different from OM. 300 mg of each preparation and biological replicates of purified OM and OMVp proteins were digested with trypsin. The resulting peptides were enriched, then analyzed via liquid chromatography coupled to tandem mass spectrometry (LC-MS/MS) as explained in Materials and Methods, followed by protein identification with Mascot search engine using the Uniprot database. Volcano plot shows OM and OMVp protein populations. Red labels indicate the proteins with the highest OMVp enrichment in comparison to OM.

**Figure 3.**
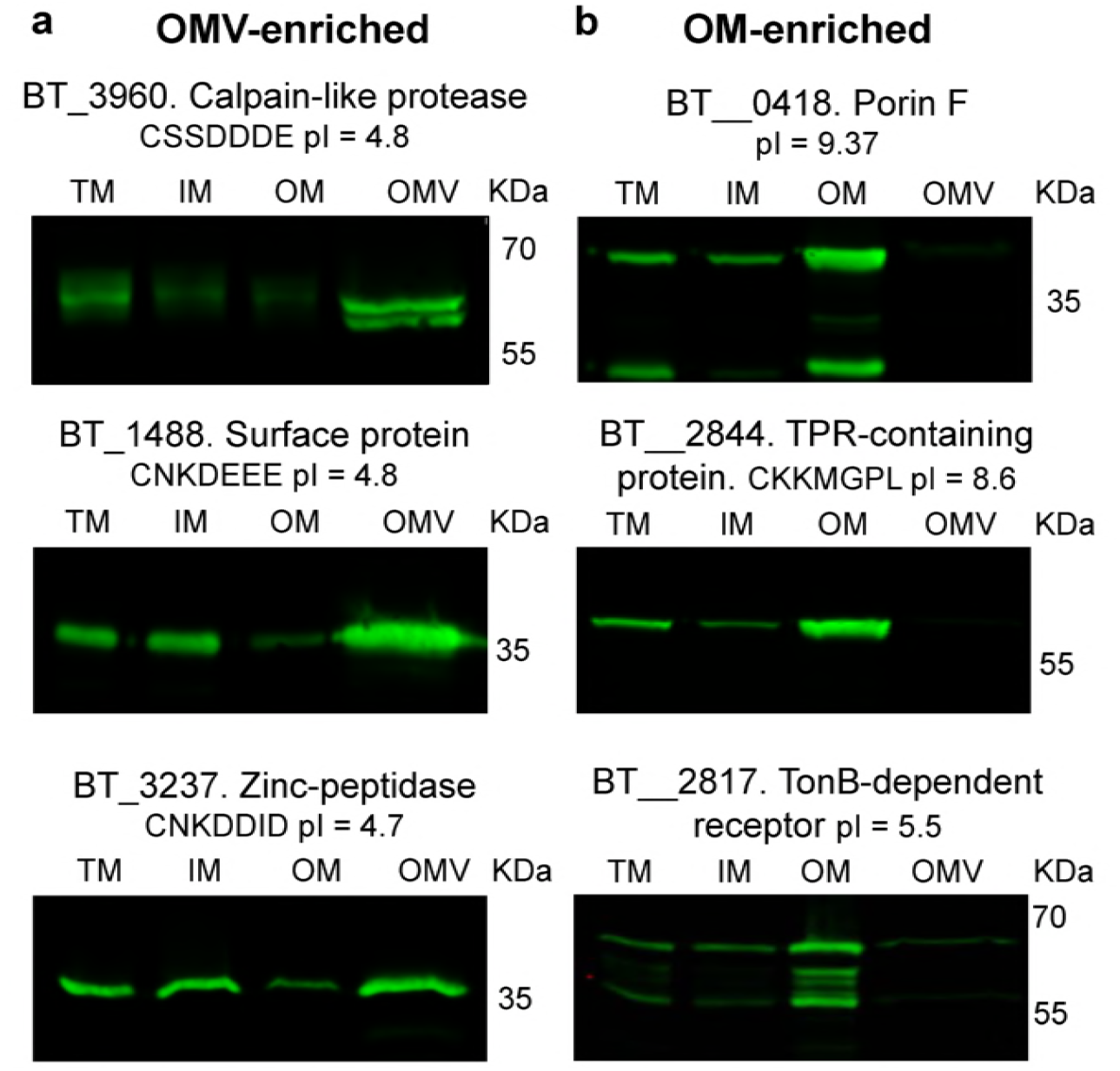
Validation of OMV‐ and OM-enriched proteins. Candidate ORFs that encode proteins identified in our MS analysis as OMV‐ or OM-enriched were cloned into pFD340 with a C-terminal 6xHis tag. Constructs were introduced into *B. theta* by conjugation, generated strains were grown in TYG media and fractions were prepared. Ten micrograms of each fraction were run on 12% SDS-PAGE and analyzed by Western Blot using anti-His polyclonal antibodies. (a) OMV-enriched proteins; (b) OM-enriched proteins. The isoelectric point as well as the residues following lipoprotein attachment Cysteine (for lipoproteins) are indicated below the protein name.

**Table 1.** Top 20 most OMVp-enriched proteins, including protein name and enrichment value. LFQ corresponds to label-free quantification values.

| Gene name | Protein name | log2 LFQ<br>(OMVp/OM) |
| --- | --- | --- |
| BT_3382 | Putative GH10/39 family protein | 13.70 |
| BT_1488 | Putative cysteine protease | 11.36 |
| BT_3960 | Putative calpain-like protease | 11.07 |
| BT_3381 | Putative GH10/39 family protein | 10.48 |
| BT_3383 | Putative GH10/39 family protein | 10.29 |
| BT_3148 | Putative adhesin | 10.12 |
| BT_2317 | Putative fimbrial protein | 9.61 |
| BT_p548229 | Putative fimbrial protein | 9.21 |
| BT_3147 | Putative adhesin | 9.18 |
| BT_1792 | Putative cell surface protein | 9.05 |
| BT_3740 | Putative cell adhesion protein | 9.02 |
| BT_3630 | Putative fimbrial protein | 8.78 |
| BT_2479 | Putative imelysin-like protein | 8.36 |
| BT_1826 | Putative GH31-like family protein | 8.29 |
| BT_3067 | Uncharacterized protein | 8.09 |
| BT_3066 | Uncharacterized protein | 7.84 |
| BT_4606 | Uncharacterized protein | 7.83 |
| BT_2658 | Putative cell adhesion protein | 7.77 |
| BT_1063 | Putative cell adhesion protein | 7.72 |
| BT_0166 | Putative thioredoxin | 7.69 |
| BT_2989 | Uncharacterized protein | 7.63 |

### Common features of OMV-enriched lipoproteins

Consistent with our previous results, we found that lipoproteins enriched in OMVs are acidic, with an average isoelectric point of 4.86 (5). We also determined that OMV-enriched lipoproteins possess a negatively charged-rich amino acid motif S(D/E)_3_ adjacent to the Cysteine residue required for lipoprotein attachment (Figure 4, Table S2). A similar motif (K-(D/E)2 or Q-A-(D/E)2) has recently been described in the oral pathogen *Capnocytophaga canimorsus*, a member of the Bacteroidetes phylum. The report showed that this motif functions as a lipoprotein export signal (LES) required for surface exposure of OM lipoproteins (28). We found that many OMV-enriched *B. theta* lipoproteins are putative protein and sugar hydrolases, required for the breakdown of complex nutritional sources. The presence of a LES on OMV proteins is consistent with their annotated functions, as they are required to face the extracellular milieu to access their cognate substrates. Conversely, OM-enriched lipoproteins like BT_2844, do not carry a LES motif. Instead, lipoproteins preferentially retained at the OM have lower sequence conservation and lower frequency of negatively-charged amino acids along the residues adjacent to the lipoprotein attachment cysteine (Figure 4b, Table S2).

**Figure 4.**
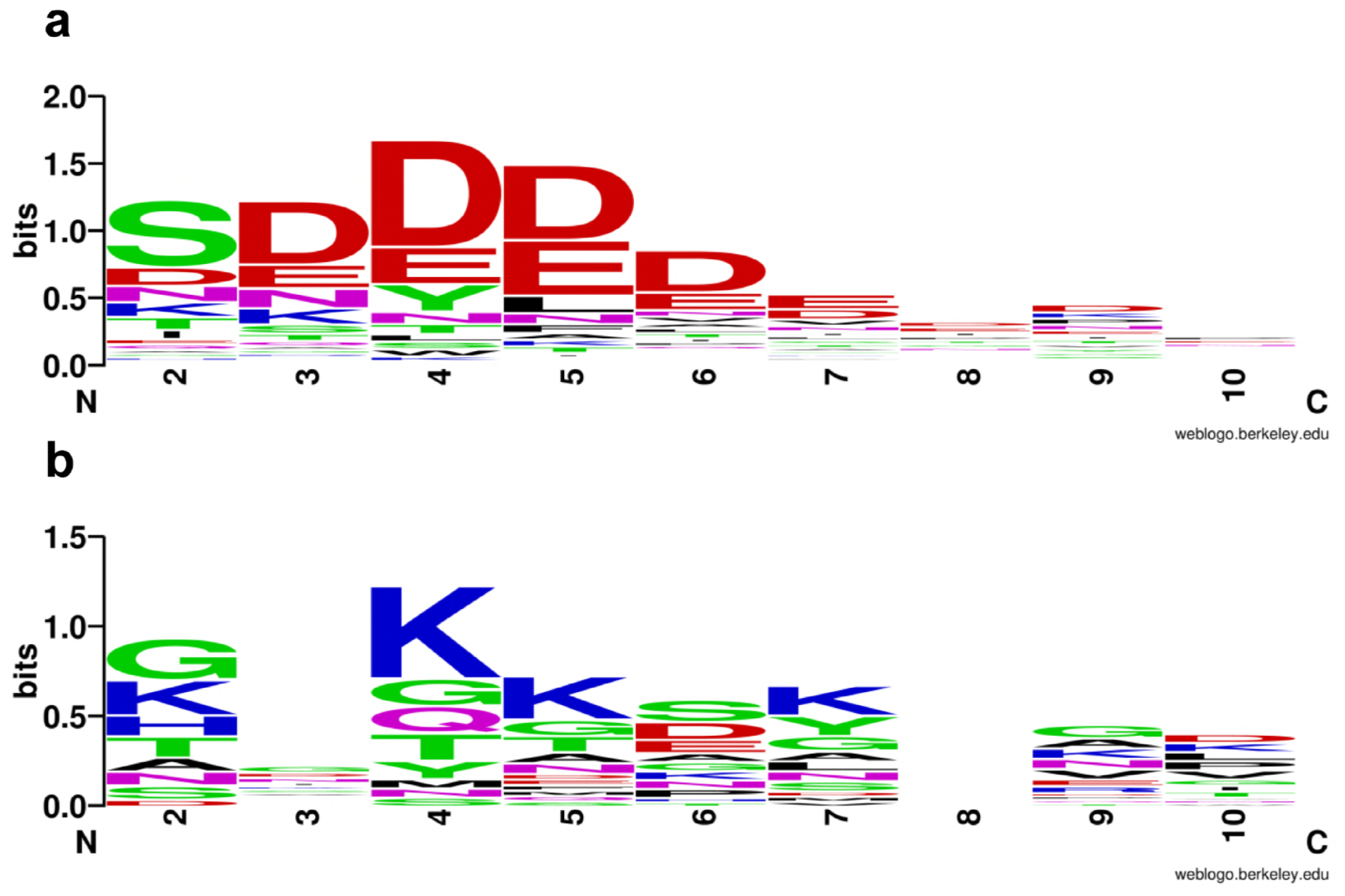
OMV-enriched proteins show a conserved N-terminal LES motif. The top 91 OMV-enriched proteins (a) and top 24 OM-enriched proteins (b) were aligned using the lipoprotein attachment cysteine as the +1 position (not shown in the logo), plus the 9 C-term contiguous residues. Lipoprotein export sequence (LES) consensus was generated using WebLogo (https://weblogo.berkeley.edu/logo.cgi)(53).

### SusG and other Sus lipoproteins are enriched in OMV and SusC is retained at the OM

In *B. theta*, hydrolytic lipoproteins are mainly encoded in PULs. PUL and PUL-like operons account for approximately 20% of the *B. theta* genome. 36 lipoproteins encoded in PUL and PUL-like operons were found in our OMV-enriched proteins. One of the most studied lipoproteins from *B. theta* is the α-amylase SusG, encoded by the *sus* operon and essential for starch catabolism (29, 30). The *sus* operon has been shown to be induced by starch and maltooligosaccharides (31). Our MS data shows that even in non-inducing conditions, two lipoproteins encoded by the *sus* operon, SusD and SusE, are enriched in OMVs (Table S2). Given the importance of starch in the mammalian diet, we investigated the subcellular localization of the components of the Sus operon. We found that all Sus lipoproteins, in particular SusG, are enriched in the OMV. The only exception was the porin SusC, which is mostly retained at the OM (Figure 5). This result strongly suggests that hydrolytic lipoproteins can exert their biological effect not only at the level of the outer membrane but also through OMVs.

**Figure 5.**
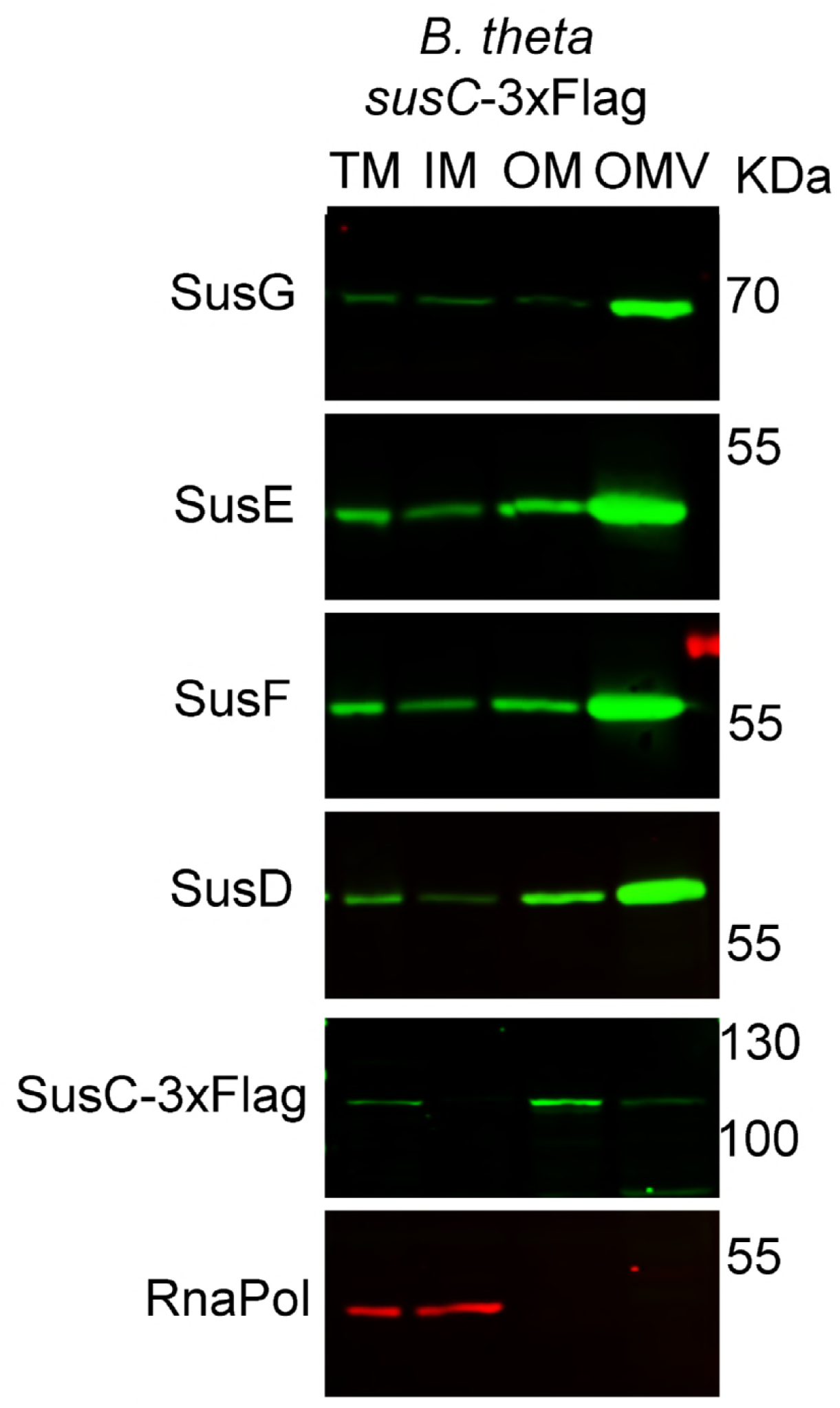
Sus lipoproteins are OMV-enriched while SusC is OM-enriched. *B. theta* cells containing a tagged genomic copy of *susC* with 3xFlag were grown in TYM media (TYG recipe with 0.5% maltose instead of glucose for induction of the *sus* operon). Fractions were prepared and analyzed by 12% SDS-PAGE and Western Blot using specific anti-Sus, anti-Flag and anti-RnaPol antibodies.

### LES is required for SusG exposure and packing into OMVs

All *B. theta* endoglycanases involved in the first step of polysaccharide breakdown, including SusG, are surface-exposed lipoproteins (29, 32). Because most OMV-enriched lipoproteins contain a LES motif, we investigated a possible association between surface exposure and packing into OMVs. SusG only contains two Asp residues following the +2 Ser residue, and therefore, for the purpose of this work, we provisionally define its LES motif as CSDD. We performed a mutational analysis on the LES motif of SusG and analyzed the localization of the protein by fractionation and Western blot (Figure 6). We cloned WT His-tagged SusG and a set of proteins carrying amino acid replacements in the LES (Figure 6a) and expressed them in the Δ*susG* strain (33). Replacement of the lipid attachment site (C23) by Ala resulted in abrogation of SusG-recruitment into OMVs. Furthermore, substitution of a single Asp residue with Ala (CSAD, CSDA, CAAD) decreased OMV-packaging (~50% relative to WT), while mutation of both Asp residues by either Ala or Lys (CSAA, CAAA, CAKK) had a more dramatic effect (~15% relative to WT) (Figure 6c). Conservative replacement of Asp by Glu (CSEE) displayed a WT-like behavior. These results indicate that the LES motif CS(D/E)_2_ is required for SusG packaging into OMVs.

**Figure 6.**
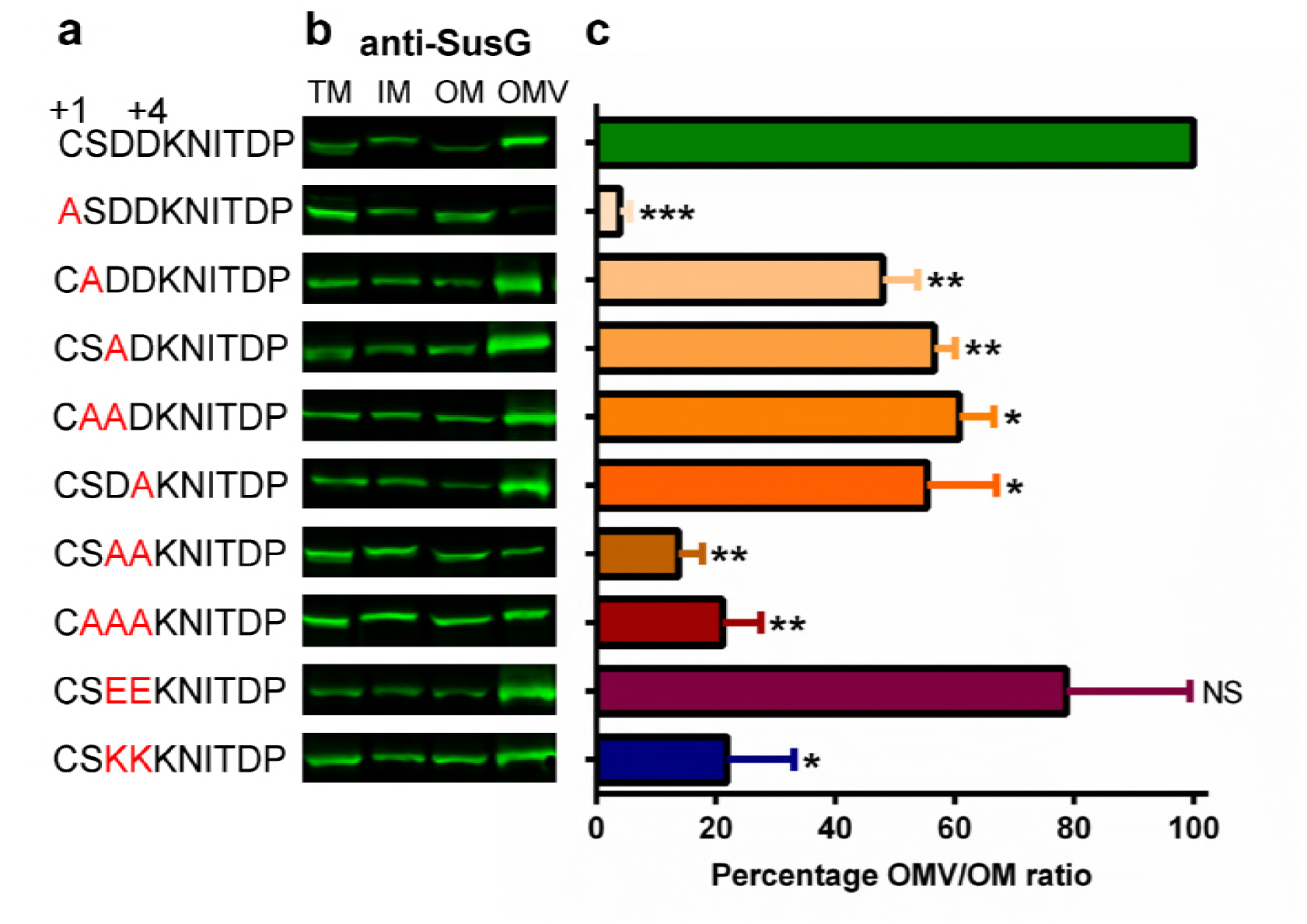
SusG LES is required for efficient packing into OMVs. (a) SusG LES point mutants were generated on pFD340/*susG*-6xHis using Quick-site directed mutagenesis. (b) Constructs were introduced in the Δ*susG* background, and generated strains were grown in minimal media with glucose as only carbon source. Fractions were prepared and analyzed by SDS-PAGE and Western Blot using anti-SusG antibodies. (c) OMV/OM ratios were calculated using the fluorescence signal values for each protein and plotted as a percentage relative to the WT OMV/OM ratio (100%). Statistical significance was determined by unpaired t-test of each LES variant in comparison to wild-type LES strain. * p-values <0.03. Experiment is representative of three biological replicates, shown are mean values with SD for two technical replicates.

The LES motif has been defined as a surface exposure tag in *C. canimorsus* (28). We used the WT LES SusG construct as well as the non-Asp variants (CSAA, CAAA) to determine whether this motif is also required for surface exposure in *B. theta*. Whole cells and OMVs from the different strains were subjected to Proteinase K (ProK) sensitivity analysis. Only the WT SusG was degraded by ProK (Figure 7). As a control, we employed a periplasmic soluble protein that localizes into the lumen of OMVs (BT_0766) and is therefore protected from ProK degradation. Taken together, these experiments confirm that LES mediates both SusG surface exposure and enrichment in OMVs.

**Figure 7.**
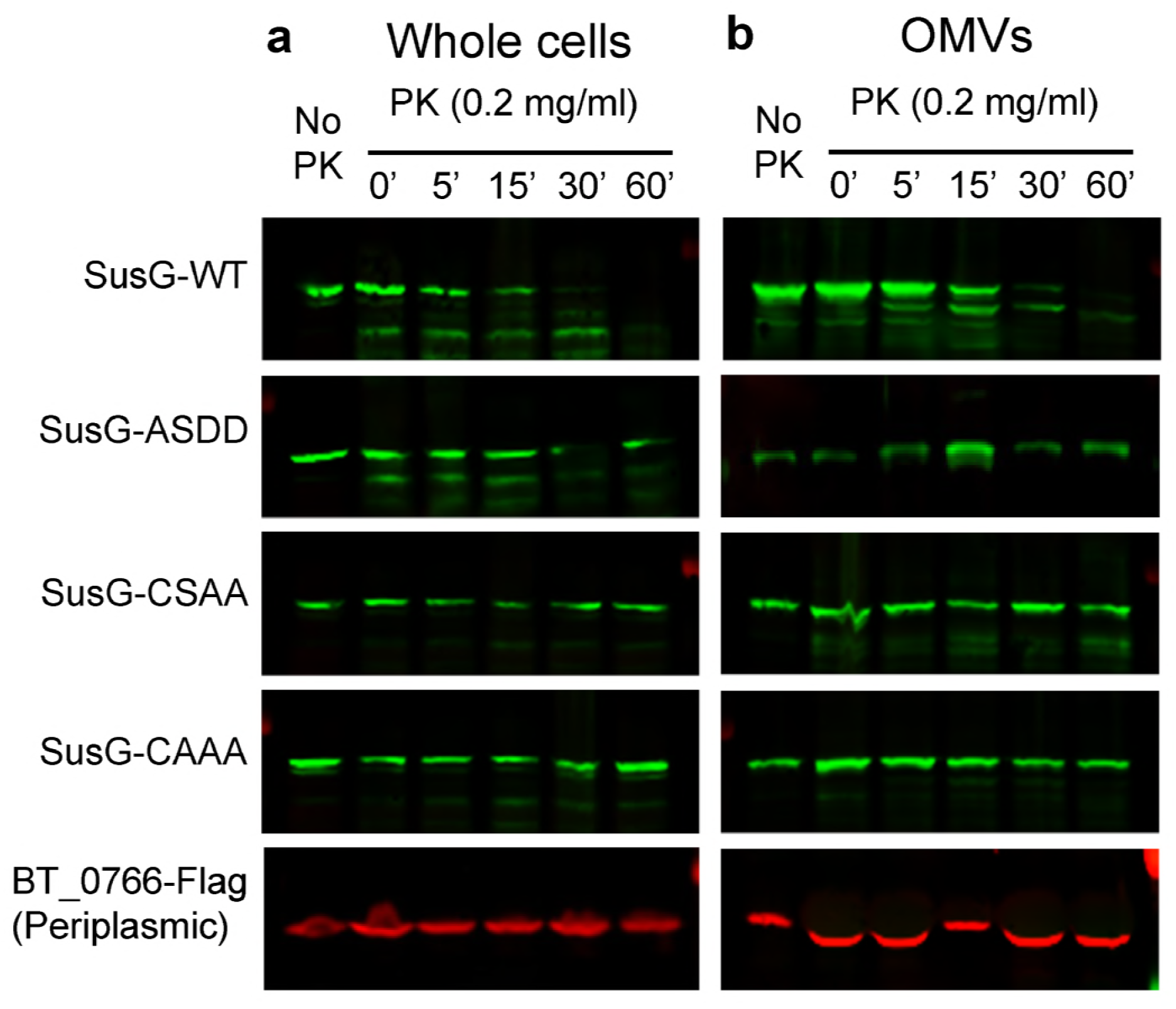
The LES is required for SusG surface exposure. WT SusG and the LES mutants were assessed in their surface exposure by proteinase K assays using whole cells (a) and OMVs (b). Proteinase K was added and incubated at 37°C; at different times, aliquots were TCA-precipitated and analyzed by SDS-PAGE and Western Blot using anti-SusG antibodies. We used a 3xFlag-tagged BT_0766, a periplasmic soluble protein found in OMVs, as an outer membrane and OMV integrity control.

### OMV containing SusG rescue Δ*susG* growth in starch

Our experiments demonstrated that SusG is packed and surface-exposed in OMVs. OMVs carrying certain glycosyl hydrolases can digest complex polysaccharides providing essential nutrients to bacteria unable to degrade these substrates (22, 23). The Δ*susG* strain is unable to grow on minimal media with starch as the sole carbon source (Figure 8). We investigated whether surface exposure of OMV-delivered SusG can rescue the Δ*susG* growth phenotype on starch. For this, we employed OMVs produced by the Δ*susG* strain expressing SusG with mutated LES (ASDD, CSAA, and CAAA). ASDD carries soluble SusG in the periplasm, while the CSAA and CAAA SusG variants face the periplasmic side of the OM (Fig. 7). Purified OMVs were added to the Δ*susG* strain in minimal media containing starch. Only WT OMVs restored growth of the Δ*susG* strain to wild-type levels (Figure 8). OMV from Δ*susG* carrying vector control (pFD340) or expressing the non-lipoprotein mutant (ASDD), or the CSAA mutant, were unable to rescue the growth of SusG, while the mutant CAAA displayed an intermediate phenotype. Altogether, our results indicate that OMVs carrying surface exposed SusG can mediate cross-feeding of other bacteria.

**Figure 8.**
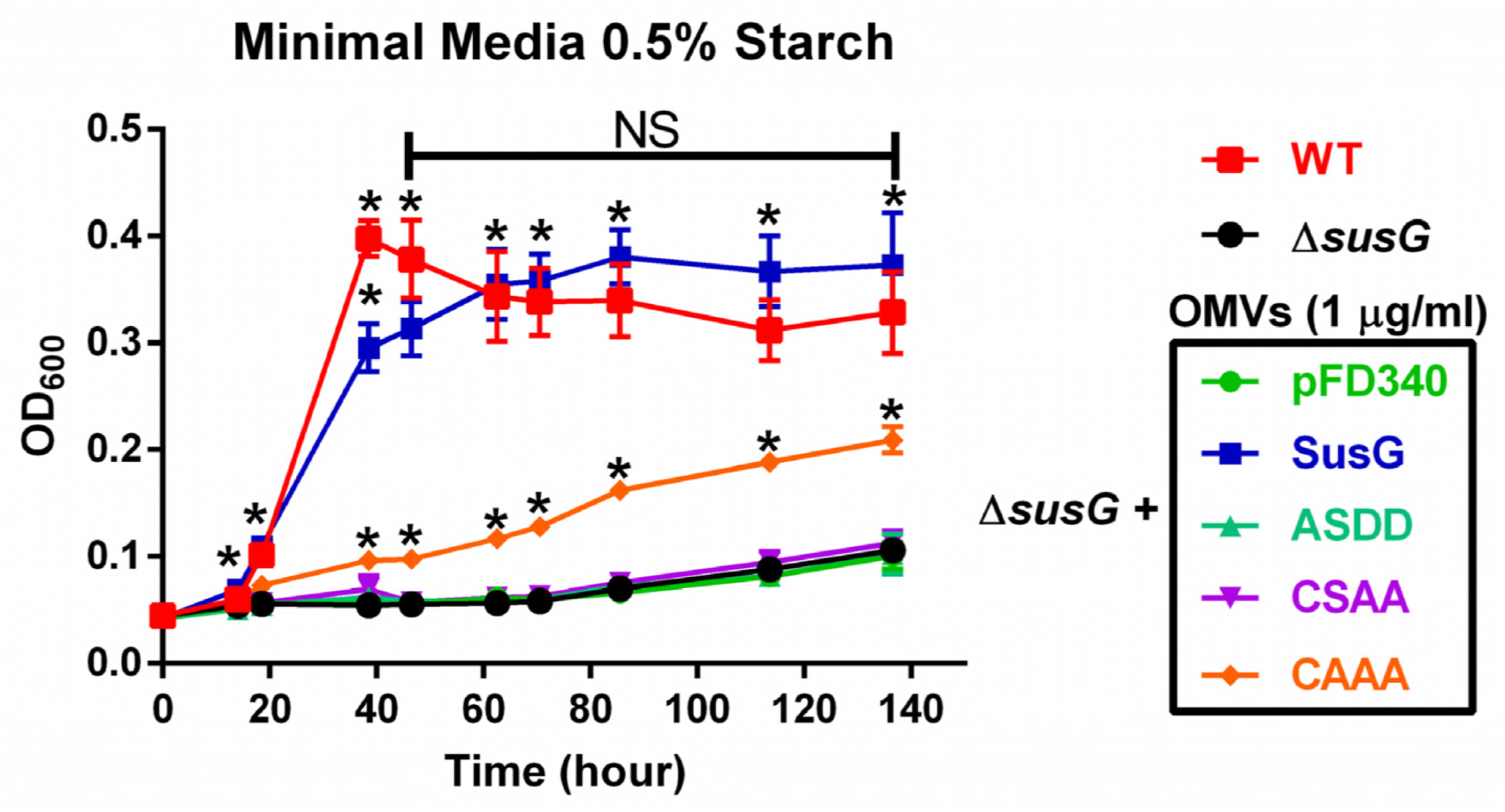
OMVs displaying WT SusG can rescue a Δ*susG* growth phenotype on starch as carbon source. WT and Δ*susG* strains were grown overnight in TYG media. Cultures were washed with minimal media (MM) without any carbon source and normalized by OD. Minimal media with 0.5% starch as only carbon source was inoculated with WT or Δ*susG* strains to final OD_600_ = 0.05. OMVs purified from the Δ*susG* strain containing different pFD340 derivatives, were added to the Δ*susG* cultures in minimal media with starch (1 μg/ml final of OMVs). Aliquots were taken at different times and OD_600_ was measured to determine growth. Statistical significance when comparing two growth curves was determined by performing one pair of unpaired t-test analysis per each time point. * p-values <0.01. Experiment is representative of three biological replicates, shown are mean values with SD for three technical replicates.

## DISCUSSION

The human gut microbiome is largely composed of species from the *Bacteroides* genus. *Bacteroides* spp are important for gut homeostasis (14, 15). They establish ecological interactions with each other and are also involved in host-commensal relationships, especially regarding the development of the immune system (7, 8, 34). Here, we show that *B. theta* produces large amounts of uniformly sized OMVs. Using an optimized methodology for the purification of different membrane fractions, we confirmed that OMV are highly enriched with lipoproteins, particularly glycosyl hydrolases. Our MS data identified the presence of a lipoprotein exposure sequence (LES) in all OMV-enriched lipoproteins. Employing the α-amylase SusG as a model, we showed that the LES is required for surface exposure and recruitment into OMVs. To our knowledge, this is the first identification of a signal involved in protein sorting into OMVs.

A handful of lipoprotein surface transport mechanisms have been described in Gram-negative bacteria (35–38). Shuttling of a group of *Neisseria meningitidis* lipoproteins to the surface is dependent on proteins Slam1 and Slam2, although the sorting mechanism has not been defined (38, 39). Moreover, the well-studied Bam system for folding of beta-barrel proteins in the outer membrane has been shown to export specific lipoproteins (36). We have not identified orthologs of Slam1 or Slam2 by sequence similarity in any *Bacteroides* genomes. The existence of LES motifs in *B. theta* lipoproteins, together with the discovery of the LES in *C. canimorsus*, suggest the existence of a conserved phylum-wide mechanism that flips specific lipoproteins towards the extracellular milieu among Bacteroidetes. Defining a functional LES in *Bacteroides* spp. constitutes the first step towards the identification of the machinery that mediates lipoprotein surface exposure in these organisms. Our experiments confirmed that the two Asp residues in position +3 and +4 respecting the lipid attachment site (+1) are essential components of SusG LES. The proposed *B. theta* LES motif CS(D/E)_2_ is similar but not identical to the (K-(D/E)_2_ or Q-A-(D/E)_2_) LES proposed for *C*. *canimorsus*. An exhaustive mutagenesis analysis of multiple proteins from several species will be required to exactly define a consensus sequence for the LES motif among Bacteroidetes.

Our results suggest that the LES has a dual role in surface exposure and recruitment of lipoproteins into OMVs (Fig 9). It is possible that the LES is involved in protein packing into OMVs through a direct interaction with an unknown component of a putative sorting machinery. Another possibility is that LES only mediates surface exposure, and flipped lipoproteins are subsequently directed for enrichment into OMVs. In the case of SusG, we have confirmed that LES is required for packing into OMVs. Future work is required to unravel the link between surface exposure and OMV packing.

**Figure 9.**
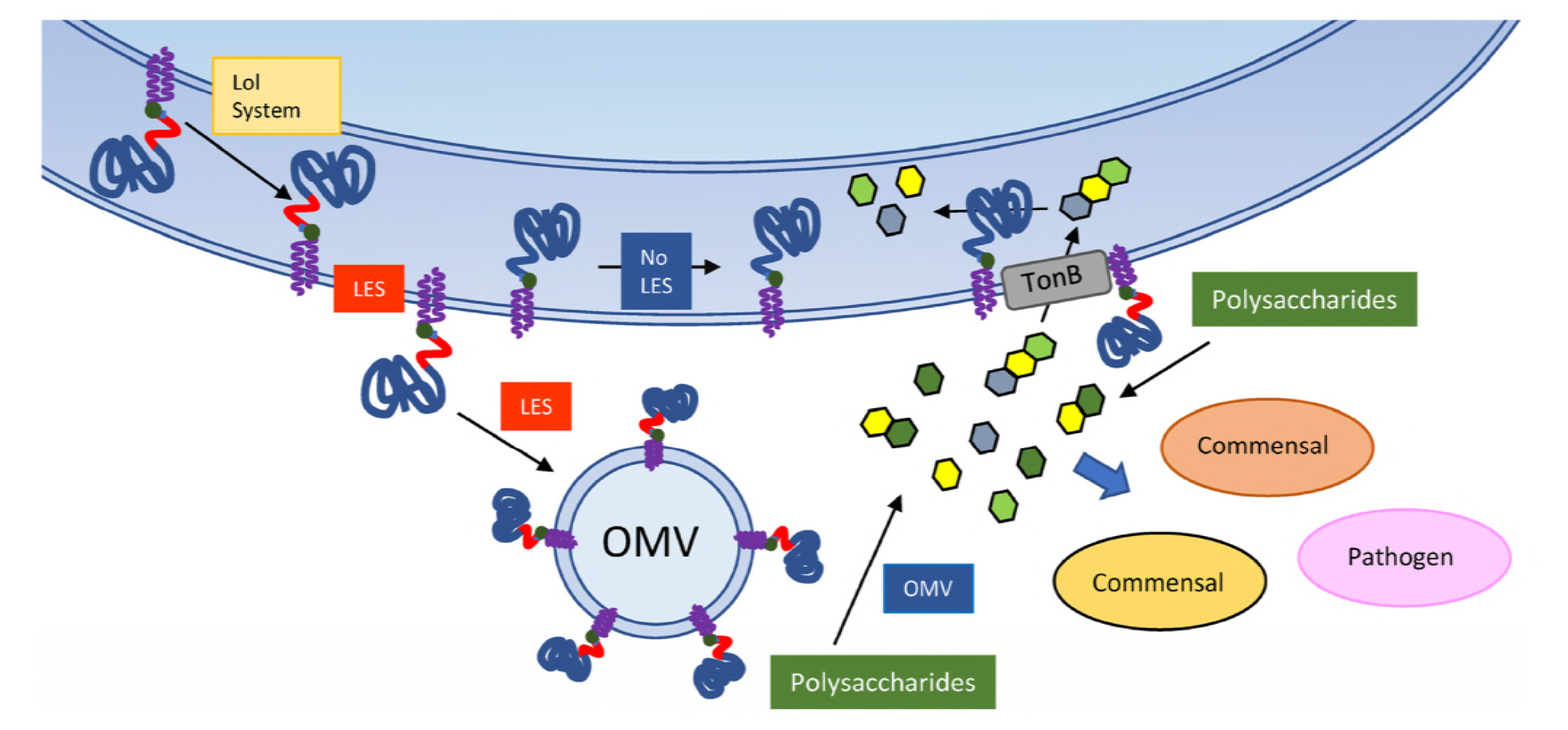
A model for SusG LES-mediated surface exposure and packing into OMV. The SusG lipoprotein is probably transported to the OM by a machinery homologous to the LOL system. Presence of a LES sequence mediates surface exposure of SusG and incorporation into OMVs. The LES could have a dual role where it is required for both surface exposure and OMV packing, or it could be involved only in surface exposure and additional information on each lipoprotein is recognized for packing into OMVs. Surface-exposed SusG in OMVs as well as in the OM hydrolyzes starch molecules into oligosaccharides that can be imported by TonB-dependent receptors by the OMV-producing cell, as well as other commensal and pathogen organisms.

In the current models for PUL systems, the hydrolytic enzymes and the oligosaccharide importers are present at the outer membrane (20, 32). However, we determined that functional glycosyl hydrolases are mainly packed into OMVs. We also found that the glycosidases present in OMVs can provide substrates to support the growth of other bacteria, in agreement with the previously proposed function of OMVs as public goods (22, 23). We propose a model in which, in a process involving the LES, the glycosidases are preferentially sorted into OMVs (Fig 9). The secreted OMVs, armed with an arsenal of hydrolases, can digest diverse dietary polysaccharides and host glycoconjugates, making the mono- and oligo-saccharides available to all members of the microbiota. Depending on the diet composition, *Bacteroides* spp can induce and package different enzyme repertoires, and therefore, members of the microbiota can act as donors or acceptors in their ecological niches. In addition, pathogens such as *Campylobacter*, *Salmonella*, and *Clostridium* can also benefit from the glycosidic activity contained in OMVs (40, 41).

We have identified a set of OMV- and OM-enriched proteins that may be employed as markers to investigate the process of OMV biogenesis through the visualization of OMV in the mammalian gut. For example, these markers would allow the differentiation *in vivo* between *bona-fide* OMV and cell-lysis derived material. Furthermore, the identification of LES as a signal for surface exposure and transport into OMVs constitutes a starting point for the design of novel probiotic interventions in animal and human health. In the future, the LES of SusG or other OMV-proteins could be employed to engineer *Bacteroides* strains to secrete OMVs packed with medically-relevant proteins in the mammalian gut.

## MATERIALS AND METHODS

### Bacterial strains and growth conditions

Oligonucleotides, strains and plasmids are described in Table S3. *Bacteroides* strains were grown in an anaerobic chamber (Coy Laboratories) using an atmosphere of 10% H_2_, 5% CO_2_, 85% N_2_. For liquid growth, TYG, TYM (TY media supplemented with 0.5% maltose instead of 0.5% glucose), or minimal media supplemented with 0.5% glucose or 0.5% potato starch were prepared as previously described (17, 42). BHI-Agar with 10% defibrinated horse blood was used as solid media. Antibiotics were used as indicated; Amp, 100 μg/ml; Erm, 25 μg/ml; BrdU, 200 μg/ml.

### Construction of plasmids, mutagenesis and generation of *susC*-3xFlag strain

Constructs for over-expression were built on pFD340 (43). Inserts obtained by PCR were digested with restriction enzymes, purified and ligated onto digested vector pFD340 as previously reported. For cloning of BT_0766-Flag into pFD340, purified PCR inserts were integrated into PCR-amplified pFD340 using InFusion Cloning Kit (Clontech). Mutagenesis of pFD340/*susG*-6xHis was carried out by inverse PCR with Pfu Turbo (Agilent) using overlapping oligos carrying the mismatch required as described elsewhere. Generation of a *susC*-3xFlag strain was carried out using the *B. theta* Δ*tdk* strategy as previously described (44). Briefly, 1000 bp upstream and downstream fragments of a C-terminal *susC*-3xFlag translational fusion were cloned into pExchange-*tdk*. Constructs were conjugated into *B. theta* Δ*tdk* cells using previously transformed *E. coli* S17-1 λpir as a donor, and strain plating and selection was performed as previously described (44).

### OMV preparations

Outer membrane vesicles were purified by ultracentrifugation of filtered spent media as previously described by our group (5). For MS analysis, OMV preparations were resuspended in 50 mM HEPES pH 7.4 and N-lauroyl sarcosine was added to 1% final concentration in 1.5 ml polyallomer tubes (Beckman Coulter). Samples were incubated with gentle rocking for 1 hour at room temperature (RT) and ultracentrifuged at 100000 x g for 2 hours at RT. Supernatants were recovered (OMV-S) and pellets (OMV-P) were resuspended in 50 mM HEPES pH 7.4. Protein content was quantified using DC Protein Assay Kit (Bio-Rad). Fractions were lyophilized for MS analysis.

### Membrane preparations

Total membrane preparations were performed by cell lysis and ultracentrifugation as previously described(5). For separation of IM and OM, total membranes were resuspended in 50 mM HEPES pH 7.4 using a 2 ml glass tissue grinder with a PTFE pestle (VWR), and N-lauroyl sarcosine was added to 1% final concentration in 1.5 ml polyallomer tubes (Beckman Coulter). Samples were incubated with gentle rocking for 1 hour at room temperature (RT) and ultracentrifuged at 100000 x g for 2 h at RT. Supernatants were recovered (IM) and pellets were resuspended in 50 mM HEPES pH 7.4. Protein content was quantified using DC Protein Assay Kit (Bio-Rad). Fractions were lyophilized for MS analysis.

### Mass spectrometry analysis

#### Protein clean up and In-solution digestion

Lyophilized protein preparations were solubilized in lysis buffer (4% SDS, 10mM, 100mM Tris pH8.5) by boiling for 10 min and the protein content assess by BCA protein assay according to the manufactures instruction. 300 μg of each preparation and biological replicate were adjusted to a volume of 200 μl and precipitated overnight using 800 μl of ice-cold acetone (1:4 v/v). Sample were spun down at 16,000G for 10min at 0°C to the resulting protein precipitate and acetone remove. Residue acetone was driven off at 70°C for 5 min. Protein precipitants were resuspended in 6 M urea, 2 M thiourea, 40 mM NH_4_HCO_3_ and reduced / alkylated prior to digestion with Lys-C (1/200 w/w) then trypsin (1/50 w/w) overnight as previously described (45). Digested samples were acidified to a final concentration of 0.5% formic acid and desalted with using C_18_ stage tips (46, 47)

#### LFQ based quantitative proteome LC-MS

Purified peptides prepared were resuspend in Buffer A* and separated using a two-column chromatography set up composed of a PepMap100 C18 20 mm x 75 μm trap and a PepMap C18 500 mm x 75 μm analytical column (Thermo Fisher Scientific). Samples were concentrated onto the trap column at 5 μL/min for 5 minutes and infused into an Orbitrap Q-exactive^™^ plus Mass Spectrometer (Thermo Fisher Scientific) at 300 nl/minute via the analytical column using a Dionex Ultimate 3000 UPLC (Thermo Fisher Scientific). 125-minute gradients were run altering the buffer composition from 1% buffer B to 28% B over 95 minutes, then from 28% B to 40% B over 10 minutes, then from 40% B to 100% B over 2 minutes, the composition was held at 100% B for 3 minutes, and then dropped to 3% B over 5 minutes and held at 3% B for another 10 minutes. The Q-exactive^™^ Mass Spectrometer was operated in a data-dependent mode automatically switching between the acquisition of a single Orbitrap MS scan (60,000 resolution) and 15 MS-MS scans (Orbitrap HCD, 35,000 resolution maximum fill time 110 ms and AGC 2*10^5^).

#### Mass spectrometry data analysis

Identification and LFQ analysis and was accomplished using MaxQuant (v1.5.3.1) (48). Searches were performed against the *B. thetaiotaomicron* (strain ATCC 29148 / VPI-5482) proteome (Uniprot proteome id UP000001414, downloaded 20-05-2017, 4,782 entries) with carbamidomethylation of cysteine set as a fixed modification and the variable modifications of oxidation of methionine and acetylation of protein N-termini. Searches were performed with trypsin cleavage specificity allowing 2 miscleavage events with a maximum false discovery rate (FDR) of 1.0% set for protein and peptide identifications. To enhance the identification of peptides between samples the Match Between Runs option was enabled with a precursor match window set to 2 minutes and an alignment window of 10 minutes. For label-free quantitation, the MaxLFQ option within Maxquant (49) was enabled in addition to the re-quantification module. The resulting protein group output was processed within the Perseus (v1.4.0.6) (50) analysis environment to remove reverse matches and common protein contaminates prior. For LFQ comparisons missing values were imputed using Perseus. Visualization was done using Perseus and R. Predicted localization and topology analysis for proteins identified by MS was performed using LipoP and TOPCONS (51, 52).

### SDS-PAGE and OMV/OM ratio determination

Membrane and OMV fractions were analyzed by standard 10-12% Tris-Glycine SDS-PAGE as described elsewhere. Briefly, 10 μg of each fraction (TM/IM/OM/OMV) were loaded onto SDS-PAGE gels, transferred onto nitrocellulose membranes and Western Blots were performed using the LI-COR system. Membranes were blocked using TBS-based Odyssey blocking solution (LI-COR). Primary antibodies used in this study were rabbit polyclonal anti-His (Thermo Fisher), mouse monoclonal anti-Flag M2 (Sigma), mouse monoclonal anti-*E. coli* RNApol subunit alpha (Biolegend) and mouse polyclonal anti-SusD/E/F/G (Nicole Koropatkin/Eric Martens, University of Michigan). Secondary antibodies used were IR-Dye anti-rabbit 780 and IR-Dye anti-mouse 680 antibodies (LI-COR). Imaging was performed using an Odyssey CLx scanner (LI-COR).

For validation of MS data, duplicate SDS-PAGE gels were stained by Coomassie Blue as described elsewhere and gel images were acquired to determine fraction quality and relative abundance (Fig. S2 and S3). For OMV/OM determination of SusG-6xHis experiments, cells were grown in minimal media with glucose and fraction were prepared as described. After transferring SDS-PAGE gels, nitrocellulose membranes were incubated with REVERT Total Protein Stain as described by the manufacturer (LI-COR) and imaged immediately at 680 nm (Fig. S4). After imaging, Western Blots using SusG antibodies were carried out and membranes were scanned at 780 nm. Total intensity values were calculated for each lane using the Odyssey scanner software (Image Studio, LI-COR). Total intensity values for each fraction, determined by scanning each lane of the REVERT stain image, were used to relativize each SusG signal. Relativized OMV and OM SusG fluorescence signals were used to calculate an OMV/OM SusG ratio. Such ratio was considered 100% for the WT LES SusG construct. Statistical significance between different OMV/OM ratios was determined by unpaired t-test for each pair of SusG LES variants.

### Proteinase K assays

Strains from *B. theta* were grown in minimal media with glucose and cells were washed with PBS and normalized to 9 OD_600_/ml. 540 μl of PBS and 10 μl of a Proteinase K (ProK) solution (20 mg/ml) were added to 450 μl of the cell suspension. Tubes were incubated at 37°C and 200 μl aliquots were removed at different time points and precipitated using trichloroacetic acid (TCA, 20% v/w final concentration). Precipitated aliquots were washed twice with acetone, and pellets were resuspended with Laemmli buffer for Western Blot analysis. A non-treated control of the cell suspension was incubated for the longest time point of the experiment and TCA-precipitated as described. A similar procedure was followed for OMV ProK treatments, 450 μl of purified OMVs were treated in the same conditions as whole cells.

### Growth curves and OMV complementation

For growth curves, wild-type or Δ*susG* strains were grown overnight in TYG media. Cultures were washed with minimal media (MM) without any carbon source and normalized by OD_600_. Minimal media with 0.5% starch as only carbon source was inoculated with WT or Δ*susG* strains to final OD_600_ = 0.05. OMVs were purified from Δ*susG* strains containing different pFD340/*susG*-6xHis derivatives grown in minimal media with glucose, and were added to the Δ*susG* cultures in minimal media with starch (1 μg/ml final OMV concentration). Aliquots were taken at different times and OD_600_ was measured to determine growth. Statistical significance when comparing two growth curves was determined by performing one pair of unpaired t-test analysis per each time point.

### Transmission electron microscopy

For negative staining and analysis by transmission electron microscopy, bacterial suspensions in PBS were allowed to absorb onto freshly glow-discharged Formvar/carbon-coated copper grids for 10 min. Grids were washed in dH_2_O and stained with 1% aqueous uranyl acetate (Ted Pella, Inc., Redding, CA) for 1 min. Excess liquid was gently wicked off, and grids were allowed to air dry. Samples were viewed on a JEOL 1200EX transmission electron microscope (JEOL United States, Peabody, MA) equipped with an AMT 8-megapixel digital camera (Advanced Microscopy Techniques, Woburn, MA).

## ACKNOWLEDGMENTS

We thank our colleagues from the Feldman laboratory for critical reading of the manuscript and comments. We would also like to express our gratitude to Nicole Koropatkin and Eric Martens (University of Michigan, USA) for their generosity in providing us with the Δ*susG* strain as well as with the anti-Sus specific antibodies.

## Supplementary Figure Legends

**Figure S1. Protocol for purification and analysis of *B. theta* OMV and membrane fractions.** We performed membrane and OMV fractionation as indicated using cultures grown overnight in TYG media (a). Fractions were incubated with 1% Sarkosyl for 1 h at RT and ultacentrifuged for 2 h at RT. Fractions were resuspended in 50 mM HEPES pH 7.4. Aliquots from each fraction were analyzed by SDS-PAGE and Coomassie Blue staining (b). Fractions labelled as OM (green) and OMV-P (blue) are the most relevant fractions for comparison by MS.

**Figure S2. Total protein profiles of membrane and OMV fractions from OMV-enriched protein validations.** We performed membrane and OMV fractionation as indicated using cultures grown overnight in TYG media. Duplicate SDS-PAGE gels loaded with 10 μg of each protein fraction were analyzed in their total protein profiles by Coomassie Blue stain. Provided are representative gels used for two OMV-enriched proteins BT_3960 and BT_1488

**Figure S3. Total protein profiles of membrane and OMV fractions from OM-enriched protein validations.** We performed membrane and OMV fractionation as indicated using cultures grown overnight in TYG media. Duplicate SDS-PAGE gels loaded with 10 μg of each protein fraction were analyzed in their total protein profiles by Coomassie Blue stain. Provided are representative gels used for two OM-enriched proteins BT_0418 and BT_2844.

**Figure S4. Total protein stain analysis of fractions for OMV/OM ratio determination.** We performed membrane and OMV fractionation as indicated using cultures grown overnight in minimal media with glucose. SDS-PAGE gels loaded with 10 μg of each protein fraction were transferred onto nitrocellulose membranes and subjected to REVERT total protein stain. Membranes were imaged immediately and Western Blot analysis was carried out using anti-SusG antibodies as described in Materials and Methods. Gel shown is representative of all the variants of SusG LES that were assayed for OMV/OM ratio determinations.

**Table S1. Complete MS dataset.**

**Table S2. OMV- and OM-enriched proteins.**

**Table S3. Oligonucleotides, strains and plasmids.**

